# A Telomere-to-telomere genome of the hibernating fat-tailed dwarf lemur (*Cheirogaleus medius*)

**DOI:** 10.64898/2026.07.30.741870

**Authors:** Ana M. Breit, Rafaela S.C. Takeshita, Erin Ehmke, Carrie Walls, Charles Billington, Peter Larsen, Adam McLain, Christopher Faulk

**Author notes:** Corresponding author: Ana M. Breit, University of Wisconsin-Stevens Point 2100 Main Street, Stevens Point, WI, USA 54494. Author contact information: Rafaela S.C. Takeshita, P.O. Box 5190, Department of Anthropology, Kent State University, Kent, OH, USA 44242-0001, Erin Ehmke, Duke Lemur Center, Duke University Durham, NC, USA 27705, Carrie Walls, Department of Animal Science, University of Minnesota, Saint Paul, MN, USA 55108, Charles Billington, Department of Pediatrics, University of Minnesota, Saint Paul, MN, USA 55454, Peter Larsen, Department of Veterinary and Biomedical, Sciences, University of Minnesota, Saint Paul, MN, USA 55108, Adam McLain, Department of Biology and Chemistry, SUNY Polytechnic Institute, Utica, NY, USA 13502, Christopher Faulk, Department of Animal Science, University of Minnesota, Saint Paul, MN, USA 55108.

## Abstract

Madagascar’s dwarf lemurs (genus *Cheirogaleus*) are the only obligate-hibernating primates and closest relative to humans capable of hibernation. Endemic to the increasingly fragmented dry forests of Madagascar, the fat tailed dwarf lemur (*Cheirogaleus medius*) represents a unique model for understanding primate physiology and tropical hibernation. Here we present FatTail1, a highly contiguous diploid genome assembly generated from a male *C. medius* at the Duke Lemur Center using Oxford Nanopore Technologies’ PromethION sequencing. The assembly spans 2.3 Gb, with an N50 of 103Mb, L50 of 10, and a BUSCO completeness score of over 99%. In addition to a complete mitogenome, we generated allele-specific DNA methylation profiles and annotated 23,925 genes using NCBI’s EGAPX. FatTail1 exceeds the gap-free contiguity of previously published strepsirrhine genomes, representing the first telomere-to-telomere genome of a Strepsirrhine primate, and providing a foundation for future studies of primate hibernation, epigenetic regulation, and conservation genomics.

**ARTICLE SUMMARY:** Dwarf lemurs are the only primates and closest relative to humans capable of months-long hibernation, making them an important model for understanding metabolic adaptations with relevance to human physiology. Here we present FatTail1, the first telomere-to-telomere genome assembly of a strepsirrhine primate, generated from a fat-tailed dwarf lemur (*Cheirogaleus medius*) using Oxford Nanopore Technologies’ PromethION sequencing. In addition to a highly complete nuclear genome, we assembled the mitogenome, identified allele-specific DNA methylation profiles and annotated 23,925 genes. FatTail1 exceeds the gap-free contiguity of previously published strepsirrhine genomes and provides an improved genomic resource for studies of hibernation, comparative genomics, epigenetic regulation, evolutionary biology, and conservation of this threatened primate.

## INTRODUCTION

Unlike most heterothermic mammals capable of torpor or hibernation, dwarf lemurs (genus *Cheirogaleus* Saint-Hilaire 1812) are primates and therefore share many physiological and genomic characteristics with humans[1,2]. Additionally, dwarf lemurs undergo obligate, seasonal hibernation in the subtropics, contending with relatively high ambient temperatures, unlike their temperate hibernating counterparts[1,3]. Like temperate hibernators, dwarf lemurs become hypometabolic, reduce their body temperature to ambient temperature, and reduce their heart rate and respiratory rate[4]. Although hibernation entails long periods of inactivity, hibernators do not experience the associated costs of inactivity, including severe muscle atrophy, bone loss, and insulin resistance[5–9]. As the closest relatives to humans able to hibernate, understanding the mechanisms of warm hibernation in a primate relative has implications for human medicine and the possibility of synthetic torpor.

Recent research has begun to characterize the molecular underpinnings of hibernation, including gene expression[10–13], DNA methylation[14–18], metabolism[10,19–21], immune response[22–26], and cellular stress[13,27,28]. However, many analyses are limited by the quality of available reference genomes. Therefore, having a high-quality reference genome is critical for characterizing the genetic mechanisms underlying the hibernation phenotype.

In addition to its importance from a hibernation physiology perspective, *C. medius* is of critical conservation concern. Populations of this species, like all dwarf lemur species, are almost universally in decline due to a combination of anthropogenic deforestation (slash-and-burn agriculture) and hunting[29]. Fat-tailed dwarf lemurs are currently listed as “Vulnerable” (VU) by the IUCN Red List of Threatened Species[30]. Due to understudied populations, there is ongoing debate regarding further delineation of populations within the genus and species. Therefore, genomic resources can support conservation efforts by improving estimates of genetic diversity, inbreeding, demographic history, and adaptive potential[31,32]. High quality reference genomes can also answer outstanding questions about species demography, population decline, and subspecies designations[33–35] and are a crucial tool for conservation genomics and population management of threatened species.

While several previous genome builds are available for *C. medius*, all are highly fragmented and are below the current standard of near-Telomere-to-telomere (T2T) quality that can be achieved with longread technology[33,36]. Here we present FatTail1, a phased diploid assembly for a male fat tailed dwarf lemur from the Duke Lemur Center (DLC). To our knowledge, no previous strepsirrhine genome has reached comparable levels of gap-free contiguity. Our assembly provides a foundation for future studies of hibernation physiology, aging, epigenetic regulation, and conservation genomics.

## METHODS

### DNA Sample

An 18-year-old male *Cheirogaleus medius* named Tanager, residing at the DLC (Durham, NC, USA), was chosen for genome sequencing and genome assembly (Figure 1). A second lemur, Ostrich (male, 4 years old at time of blood draw), was chosen for additional sequencing and treated similarly. Whole blood was collected post-mortem (Tanager) and during routine research sample banking (Ostrich), shipped frozen, then thawed and combined with 2 volumes of DNA Shield (Zymo Research). Genomic DNA was extracted using a Qiagen MagAttract kit (cat. 67563 Germantown, MD, USA) yielding 3 μg of DNA from 200 μL of blood. DNA quality was checked using a nano-spectrophotometer (Implen N60, Munich, Germany) and visualized on a 1% gel to check for fragmentation and RNA contamination.

**Figure 1:**
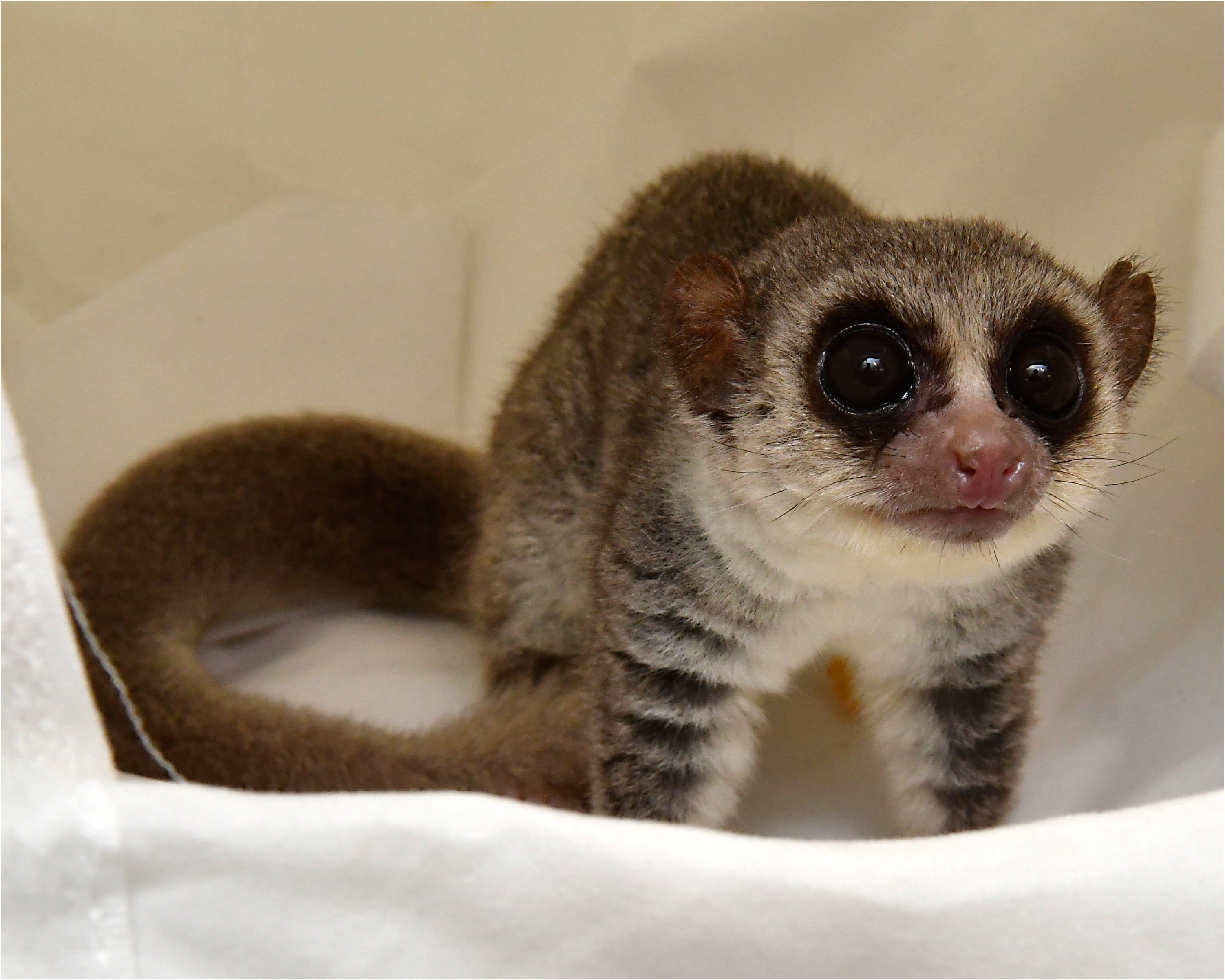
*Cheirogaleus medius* holotype photo depicting Tanager, the 18-year-old male fat-tailed dwarf lemur sampled for genome assembly. Alt Text: Tanager, an adult male fat-tailed dwarf lemur (*Cheirogaleus medius*) standing on white cloth. The animal has large forward-facing dark eyes, gray-brown fur, a pale face and underside, and a characteristically thick, fat-storing tail curled behind his body.

### Library Preparation

Libraries were prepared using the SQK-LSK114 kit (Oxford Nanopore Technologies, Oxford, UK) according to the manufacturer’s instructions. Libraries were diluted to between 20-40 ng/μL prior to sequencing.

### Sequencing

Sequencing for Tanager was performed on three R10.4.1 PromethION flow cells using Oxford Nanopore’s minKNOW software (v25.09.16)[63]. Sequencing for Ostrich was performed on two flowcells. Flowcells were run for 48 hours before library recovery, nuclease flush, reload, and run until exhaustion. Sequencing summary statistics are provided in supplementary file 1.

### Computational Methods

Detailed pipeline information including software versions is provided in Supplemental File 2. Raw sequencing data stored in pod5 files was re-basecalled using stand-alone dorado software[64] in super accuracy mode with model dna_r10.4.1_e8.2_400bps_sup@v5.2.0 with 5mC and 5hmC DNA modification detection turned on.

### Genome Assembly

Draft assemblies were generated using both Flye[65] and hifiasm assemblers[66] on the institutional cluster computing environment. Due to the size of the total read set, over 250 Gb in the case of Tanager, the assemblers required a high memory node (2 Tb) and ran for approximately 24 hours with 128-cores. Assembly was performed using the total read set as well as a HERRO corrected read set[67] for comparison. All draft assemblies were polished using dorado and results were evaluated by N50 and BUSCO. Removal of allelic haplotigs was performed with purge_dups[68]. Contigs were linked and gapfilled using the ntLinks pipeline[69]. Manual curation was performed first by comparative dotplot analysis of the draft assembly against the *Microcebus murinus* genome using the D-genies[70] website to annotate chromosomes. All contigs matching ∼1:1 in size with an *M. murinus* chromosome and having telomeres on one or both ends were kept and considered T2T. The custom telomere identification script is available in supplementary file 2. For *M. murinus* chromosomes matching to multiple *C. medius* contigs, we kept the longest contig that was T2T or had one telomere end and discarded any contigs that had no telomeric sequence and zero BUSCOs.

We performed an independent assembly of Ostrich’s genome. In several cases Ostrich’s assembly recovered complete T2T chromosomes that were fragmented in Tanager’s genome. In these cases, we used RagTag[71] to scaffold and gapfill Tanager’s fragments into complete T2T chromosomes. Remaining non-telomeric ends were BLASTed[72] for identification. Assemblies were checked for remaining sequence adapters or foreign contaminants using NCBI’s foreign contaminant screening tool[73].

### Coverage and Quality Statistics

Read and assembly summary statistics were assessed using seqkit[74] and cramino[75]. Assembly coverage was assessed using cramino and mosdepth[76]. BUSCO scores were calculated using compleasm[77] with the primate lineage. Assembly statistics were compared using quast-lg[78]. Synteny between genomes was visualized in ribbon plots using ntsynt[79]. Dotplot for publication was generated with Paf2dotplot[80].

### Variant Calling and Diploid Assembly

SNVs and short indels were called using Clair3[81] while long SVs were called using CuteSV[82], both against the alignment of reads the haploid curated draft assembly. Resultant VCFs were combined using bcftools[83]. Next, we generated a diploid assembly from the hifiasm haploid draft. Whatshap[84] phased the VCF using the haploid draft and the reads. The phased VCF was filtered for passing variants that had a Q-score greater than 20. The phased VCF was then applied to the curated assembly to create both hap1 and hap2 pseudohaplotype assemblies, each representing one parental haplotype.

Genome-wide heterozygosity was calculated as the number of variants divided by the length of the genome, excluding the sex chromosomes. ROH were assessed with *bfctools roh*, where segments ≥ 1 Mb were retained. Finally, historical N_e_ and population demographics were assessed with PSMC[85]. For each individual, we generated a diploid consensus sequence from the phased VCF files and the assembled genomes. We masked the diploid consensus with the ROH regions identified using *bcftools roh* to prevent homozygous biases. The masked genomes were converted into.psmcfa format (q ≥ 20) and analyzed using PSMC with standard atomic time intervals (4+25*2+4+6). To evaluate demographic robustness, we generated 100 bootstrap replicates by fragmenting the data with *splitfa* and randomly subsampling it via *seqtk*[86] prior to re-running the PSMC model. Final curves and confidence intervals were scaled to absolute years using a mutation rate of 1.5 x 10^-8^ and a 4-year generation time.

### DNA Methylation

Dorado called DNA methylation (5mC) and hydroxymethylation (5mC) at cytosines, and we restricted analysis to 5mC at CpG sites as the predominant modification present in mammalian blood. Phased reads were separated by parental haplotype based on shared variants. Differentially methylated regions (DMRs) were detected using the DSS R package[87]. These represent regions with greater than 65% difference in average methylation between parental haplotypes. Regions were visualized using Methylartist[88].

### Repeat Identification

Repetitive DNA masking and classification was performed with RepeatMasker[89] (V4.2.3) with the primate lineage. Annotation using existing repeat classes, families, and subfamilies was elected based on good representation of primate elements in the Dfam (V3.9) open-source repeat library[90]. To ensure consistency, we used the same pipeline to repeatmask other *Lorisidae* genomes for comparison with human as outgroup. We used the following builds: Aye-aye (GCA_044048945.1_DMad_hybrid)[41], Galago (GCF_000181295.1_OtoGar3)[91], Coquerel’s sifaka (GCF_000956105.1_Pcoq_1.0)[92], ring-tailed lemur (GCF_020740605.2_mLemCat1.pri)[93], mouse lemur (GCF_040939455.1_M.murinus_Inina_mat1.0)[94], and human (GCF_009914755.1_T2T-CHM13v2.0)[95].

### Gene Annotation

NCBI’s EGAPx[96] was used to annotate the protein-coding and non-coding genes from the primary assembly. Publicly available short-read RNA-seq data was downloaded from the NCBI sequence read archive, representing transcripts from adipose tissue derived from a fat-tailed dwarf lemur (SRR1634147, and SRR1634148). BUSCO v5.7.1 was run as part of the EGAPx pipeline, however for cross-species comparison BUSCO in protein mode was run via compleasm with primates_odb12[97] against all protein annotations associated with a genome build downloaded from NCBI.

### Mitochondrial Genome

MitoHiFi[98] generated and annotated the mitochondrial genomes from the raw reads using a previously published *C. medius* mitogenome as bait (NC_065434.1). A mitogenome phylogeny was built using mafft[99] for alignment and IQ-TREE (V.2.0.7)[100] for bootstrapped maximum-likelihood tree inference. Trees were visualized using FigTree[101].

## RESULTS

### Sequencing

DNA was isolated for genome assembly from post-mortem whole blood of an 18-year-old male captive-bred fat-tailed dwarf lemur (*Cheirogaleus medius*) named Tanager, residing at the Duke Lemur Center (DLC; Durham, NC, USA); a holotype photo of this individual is provided in Figure 1. A second lemur, Ostrich, was also sequenced from remnant whole blood for comparative analysis. Genomic DNA extraction from Tanager yielded approximately 10 μg of total DNA. Tanager’s DNA generated 58 million reads and 253 Gb of sequence with a read N50 of 9.6 kb (Table 1). Ostrich’s DNA generated 15 million reads and 81 Gb of sequence with a read N50 of 12 kb.

**Table 1:** *C. medius* nanopore sequencing read summary.

|  | Tanager | Ostrich |
| --- | --- | --- |
| Number of reads | 58,344,400 | 15,375,752 |
| Number of bases | 253,275,766,090 | 81,627,386,855 |
| N50 read length (bp) | 9,625 | 12,006 |
| Longest read (bp) | 902,906 | 707,427 |
| Mean read length (bp) | 4341 | 5309 |
| Mean read quality (Q) | 20.56 | 20.22 |
| >Q20 (%) | 93.02 | 86.76 |
| >Q30 (%) | 92.84 | 86.94 |
| GC (%) | 40.58 | 40.53 |

### Genome Assembly

Our final reference genome, FatTail1, had a size of 2.3 Gb in 38 contigs with an N50 of 103 Mb and BUSCO of over 99% completeness. This assembly derived from Tanager was phased into two pseudohaplotypes (FatTail1.hap1 and FatTail1.hap2) representing each parental genome. A second assembly was created from Ostrich and similarly phased into pseudohaplotypes CmedOstrich.hap1 and CmedOstrich.hap2.

We initially generated multiple *de novo* assemblies and chose the highest-quality result for further analysis based on the number of contigs, N50, and BUSCO score (Table 2). The hifiasm run against Tanager’s total read set yielded the most contiguous draft assembly and was selected for further analysis. The raw read quality obtained from the R10.4.1 flowcells combined with the native accuracy of the basecaller yielded a draft genome of higher quality than one generated from the corrected reads due to the higher number of uncorrected reads offsetting the lower number of corrected reads. A polishing step was attempted on the draft assembly however it reduced the number of complete BUSCOs and was therefore skipped. Further improvement was made by purging allelic duplicates, linking contigs, and manual curation.

**Table 2:** *A*ssembly statistics for prior reference genomes and assembly drafts.

| Prior reference assemblies | Size (bp) | Contigs | N50 (bp) | L50 | BUSCO | BUSCO | BUSCO | BUSCO |
| --- | --- | --- | --- | --- | --- | --- | --- | --- |
|  |  |  |  |  | Single* | Duplicate | Fragment | Missing |
| C. medius_ASM808673v1 | 2,393,092,913 | 547,751 | 118,572 | 5,476 | 89.99% | 0.28% | 2.88% | 6.84% |
| C. medius_PGDP_CheMed | 2,206,259,077 | 108,634 | 43,246 | 12,969 | 64.61% | 0.35% | 14.62% | 20.37% |
| C. medius_CheMed_v1_BIUU | 2,121,890,802 | 3,231 | 48,318,266 | 14 | 88.40% | 0.74% | 6.57% | 4.28% |
| Microcebus murinus_mat1.0 | 2,350,949,499 | 124 | 105,226,355 | 10 | 97.30% | 1.72% | 0.07% | 0.91% |
| New C. medius assembly | Size (bp) | Contigs | N50 (bp) | L50 | BUSCO | BUSCO | BUSCO | BUSCO |
|  |  |  |  |  | Single* | Duplicate | Fragment | Missing |
| Initial assembly (flye) | 2,261,795,302 | 1,287 | 22,765,254 | 30 | 98.98% | 0.91% | 0.03% | 0.08% |
| Initial assembly (hifiasm) | 2,328,478,157 | 165 | 95,502,240 | 11 | 98.96% | 0.96% | 0.03% | 0.04% |
| Purged (purge_dups) | 2,319,264,537 | 58 | 95,502,240 | 11 | 99.01% | 0.91% | 0.03% | 0.04% |
| Gap-filled (ntLink) | 2,318,947,488 | 52 | 103,143,189 | 10 | 98.99% | 0.91% | 0.03% | 0.07% |
| Curated (manual) | 2,312,045,604 | 38 | 103,143,189 | 10 | 99.03% | 0.90% | 0.03% | 0.04% |
| **FatTail1.hap1 (Tanager)** | **2,312,043,398** | **38** | **103,143,337** | **10** | **99.02%** | **0.90%** | **0.03%** | **0.04%** |
| FatTail1.hap2 (Tanager) | 2,312,029,741 | 38 | 103,142,875 | 10 | 99.03% | 0.90% | 0.03% | 0.03% |
| CmedOstrich.hap1 | 2,323,375,508 | 69 | 83,368,496 | 11 | 99.07% | 0.87% | 0.03% | 0.03% |
| CmedOstrich.hap2 | 2,323,355,561 | 69 | 83,367,578 | 11 | 99.09% | 0.85% | 0.03% | 0.03% |

Three other pre-existing *C. medius* genomes are shown for comparison along with the *Microcebus murinus* genome which is the closest sequenced relative with a genome at the chromosome level. We improved contiguity, N50, and BUSCO score over all prior assemblies, and matched or exceeded the *M. murinus* genome in all metrics.

Presumptive chromosome contigs were identified by the presence of telomeric sequence at either end as well as by comparison to the *M. murinus* genome for approximate size expectations (Table 3). We assigned chromosome numbers by order of decreasing size and were able to fully assemble 30 of the 32 autosomes to the telomere-to-telomere status. Chromosome 21 is flanked by one telomere on the 3’ side and an array of 18S ribosomal gene tandem repeats on the 5’ side which prevented complete assembly. Chromosome 30 has a telomere on the 3’ side and is flanked by a genic region that maps to human chromosome 3 on the 5’ side and remains incomplete. The sex chromosomes were also unable to be fully assembled due to their inherent repetitive nature and their haploid depth. Chromosome X consists of four unplaced contigs, two with single-end telomeres and contigs two without telomeres that represent interstitial sequence. The Y chromosome was unable to be recovered.

**Table 3.** Chromosome Assignments.

| **Chromosome assignment** | **Length (bp)** | **Telomere status** | **M. murinus homolog chr** | **Notes** |
| --- | --- | --- | --- | --- |
| Chr1 | 180126235 | T2T | 1 |  |
| Chr2 | 127984047 | T2T | 2 |  |
| Chr3 | 119954436 | T2T | 6 |  |
| Chr4 | 115067048 | T2T | 3 |  |
| Chr5 | 114597010 | T2T | 5 |  |
| Chr6 | 111254677 | T2T | 4 |  |
| Chr7 | 108923041 | T2T | 7 |  |
| Chr8 | 107758789 | T2T | 8 |  |
| Chr9 | 104675812 | T2T | 9 |  |
| Chr10 | 103143337 | T2T | 11 |  |
| Chr11 | 102396628 | T2T | 10 |  |
| Chr12 | 95502352 | T2T | 12 |  |
| Chr13 | 83320344 | T2T | 13 |  |
| Chr14 | 77935101 | T2T | 15 |  |
| Chr15 | 74199232 | T2T | 14 |  |
| Chr16 | 61558165 | T2T | 18 |  |
| Chr17 | 61036215 | T2T | 17 |  |
| Chr18 | 57940411 | T2T | 16a |  |
| Chr19 | 57491611 | T2T | 19 |  |
| Chr20 | 41119609 | T2T | 21 |  |
| Chr21 | 34688312 | 3' | 20a | 5' 18S ribosomal RNA |
| Chr22 | 33505003 | T2T | 23 |  |
| Chr23 | 31171903 | T2T | 24 |  |
| Chr24 | 28636393 | T2T | 22 |  |
| Chr25 | 26200218 | T2T | 25 |  |
| Chr26 | 24054077 | T2T | 27 |  |
| Chr27 | 22215613 | T2T | 26 |  |
| Chr28 | 22054303 | T2T | 16b |  |
| Chr29 | 17539140 | T2T | 29 |  |
| Chr30 | 15832826 | 3' | 28 | 5' homolog chr 3 H. sap |
| Chr31 | 10924678 | T2T | 30 |  |
| Chr32 | 3778372 | T2T | 20b |  |
| Unplaced_1 | 3807385 | none | Unplaced |  |
| Unplaced_2 | 2960189 | none | Unplaced |  |
| ChrX_unplaced_1 | 48595375 | 5' | X | 3' homolog Xp H. sap |
| ChrX_unplaced_2 | 45011164 | none | X | Chr X unplaced |
| ChrX_unplaced_3 | 31846560 | 5' | X | 3' homolog Xq H. sap |
| ChrX_unplaced_4 | 3237787 | none | X | Chr X unplaced |

Alignment with *M. murinus* revealed large-scale, but imperfect synteny, indicating several chromosomal fusions and fissions in the intervening 25 million years since their last common ancestor. Both species as well as all Cheirogaleidae have 33 pairs of chromosomes according to previous cytogenetic studies[37,38]. Between *M. murinus* and *C. medius* most chromosomes map directly one-to-one with some internal inversions and rearrangements (Figure 2a). *M. murinus* chromosomes 16 and 20 are each represented by two separate chromosomes in our *C. medius* assembly. Conversely the smallest *M. murinus* chromosomes, 31 and 32 are fused with other chromosomes in *C. medius*. Expanded synteny representing the four lemur families with chromosomal scale scaffolds are shown in Figure 2b revealing greater genomic rearrangement with increasing evolutionary distance.

**Figure 2:**
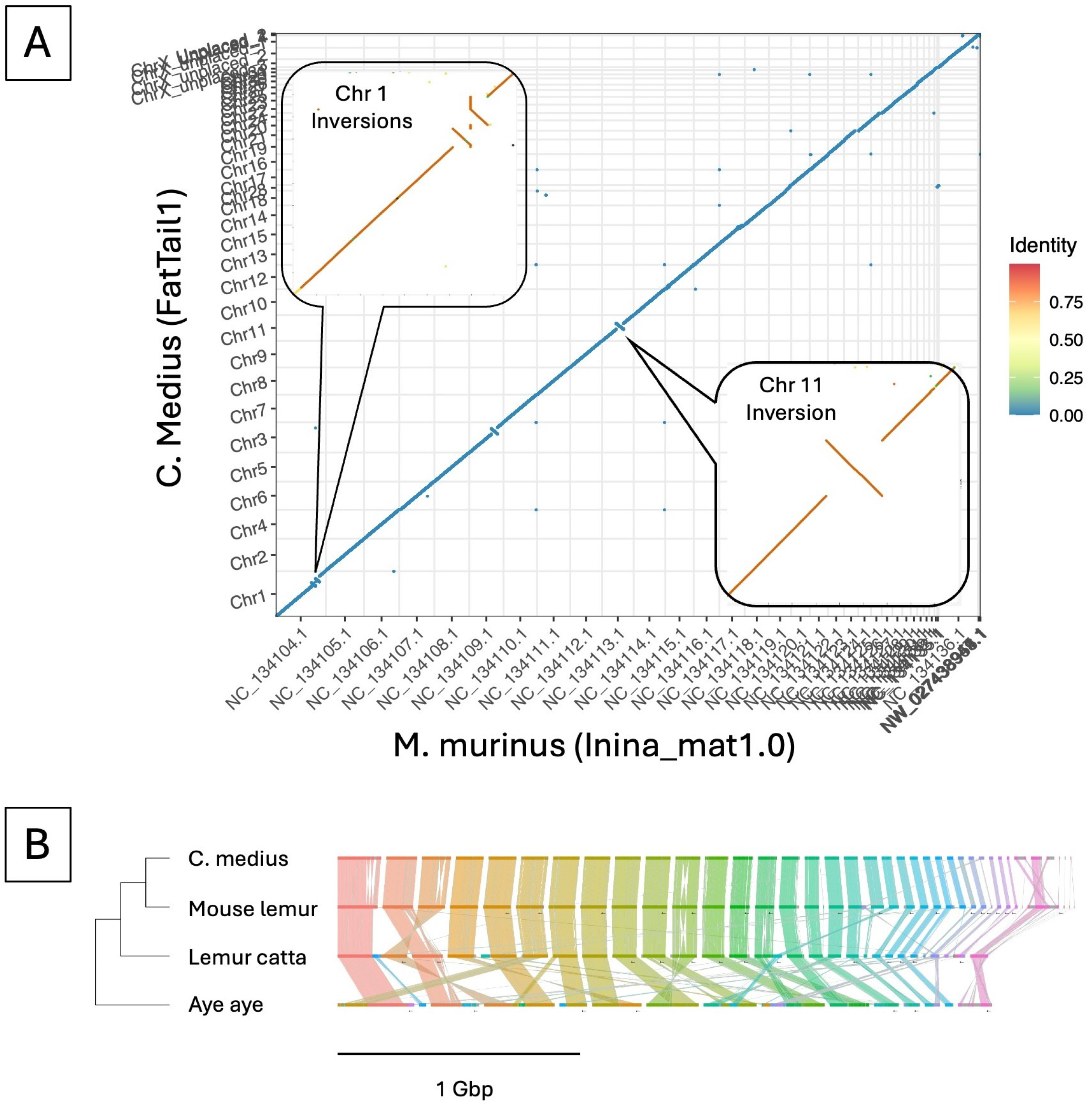
FatTail1 is a near-complete and highly contiguous genome for the fat-tailed dwarf lemur (*Cheirogaleus medius)*. (A) Genome sequence alignment of the mouse lemur (*Microcebus murinus*) reference genome (mat1.0) and FatTail1 was generated using Paf2dotplot. The alignment quality reflects gross genome collinearity present across *Cheirogaleidae*. (B) Ribbon diagram of *C. medius* chromosomes in synteny to the mouse lemur, ring-tailed lemur (Lemur catta), and Aye aye reference genomes representing greater divergence across *Lemuridae*. Color transitions represent breakpoints between contigs on the scaffold. Alt text: Comparison of the FatTail1 genome assembly with other strepsirrhine genomes. Panel A shows a whole-genome dot plot comparing the *Cheirogaleus medius* assembly to the gray mouse lemur (Microcebus murinus) reference genome. Most chromosomes exhibit strong one-to-one synteny, with two highlighted inversion regions on chromosomes 1 and 11. Panel B shows chromosome-scale synteny among *C. medius,* gray mouse lemur, ring-tailed lemur, and aye-aye genomes, illustrating overall conservation of chromosome structure across these strepsirrhine species.

The high contiguity and diploid resolution of this assembly enables simultaneous investigation of structural variation, allele-specific methylation, demographic history, and runs of homozygosity within a single endangered primate system. Currently there are chromosomal-level strepsirrhine reference genomes for the ring-tailed lemur (*Lemur catta*)[39], gray mouse lemur (*Microcebus murinus*)[40], and aye-aye (*Daubentonia madagascariensis*)[41]. Ours will be the first chromosomal-level assembly of a member of the Cheirogaleidae family, and we believe it will substantially advance comparative primate genomics.

### Repetitive DNA

Repetitive DNA, particularly interspersed transposons, are major drivers of genetic diversity in vertebrates[42]. Since primates are well-represented in transposable element (TE) databases, we classified and annotated FatTail1 repetitive DNA with RepeatMasker (Table 4). Overall, 42.54% of the assembly was classified as repetitive DNA, with a majority consisting of retroelements (35.98%) and fewer DNA elements (5.23%), simple repeats (0.96%), and low-complexity regions (0.2%) than other lemurs. Analysis of other primate assemblies with the same RepeatMasker pipeline indicated that FatTail1’s repeat complement was consistent with other primates (Figure 3). As a group, *Lemuridae* had nearly no satellite DNA detected, unlike the human genome comparator, and were depleted in simple repeats as well. In all other categories, the fat-tailed dwarf lemur was unremarkable. These results are consistent with the dominance of transposons in mammalian genomes generally.

**Figure 3:**
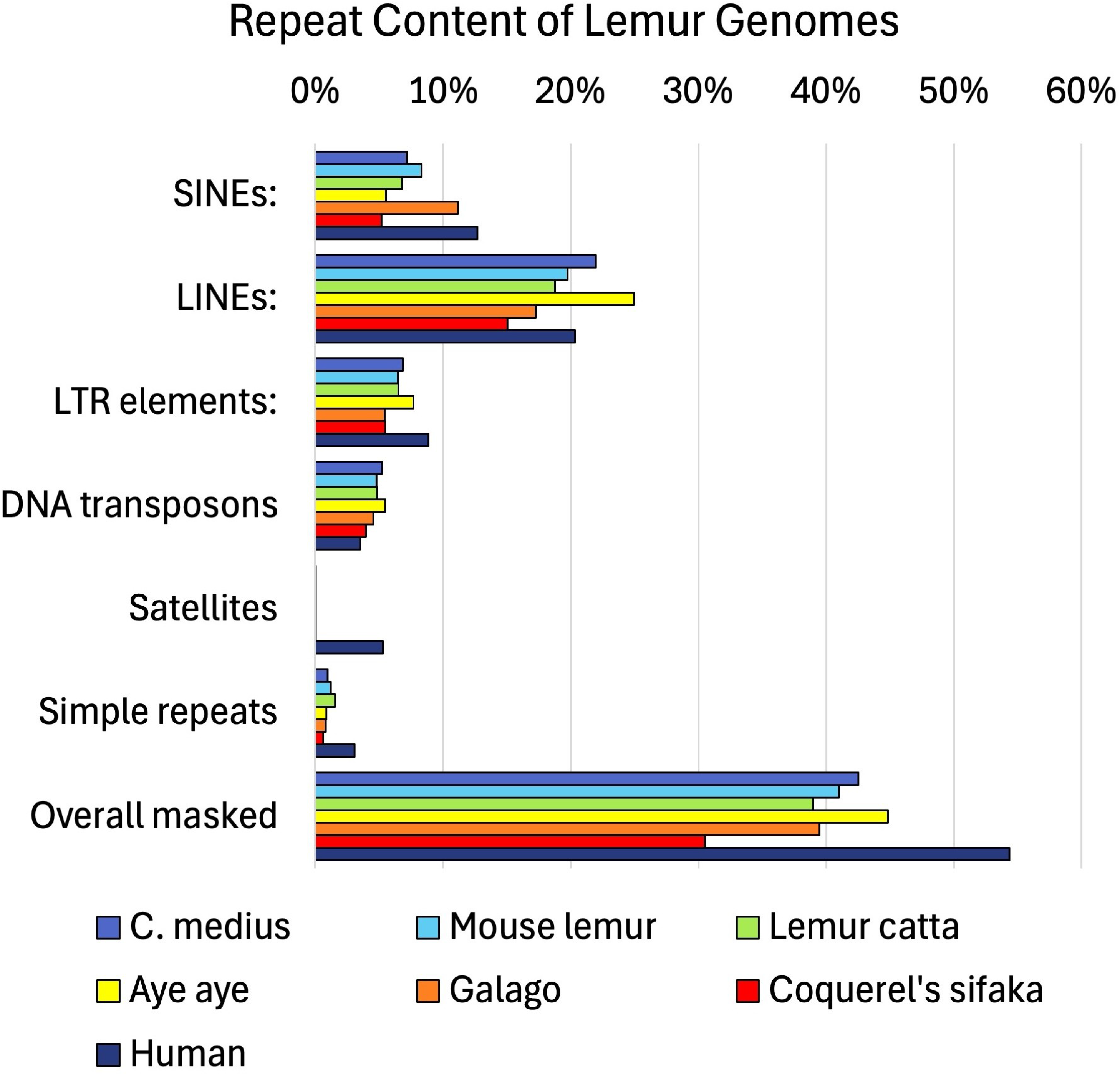
Repetitive DNA percentage. Proportion of the genome covered by repeat families for the *C. medius* genome assembly, FatTail1, as compared to reference genomes of the other listed species generated with RepeatMasker. Alt text: Horizontal bar chart comparing the proportion of repetitive DNA classes among seven primate genomes: *Cheirogaleus medius, the Aye aye, Human, Mouse Lemur, Galago, Lemur catta, and coquerel’s sifaka.* Categories include SINEs, LINEs, LTR elements, DNA transposons, satellites, simple repeats, and toverallmasked sequence. The fat-tailed dwarf lemur displays repetitive DNA content like other strepsirrhine primates, with LINE elements comprising the largest repeat class and an overall repeat content of approximately 43% of the genome.

**Table 4:**
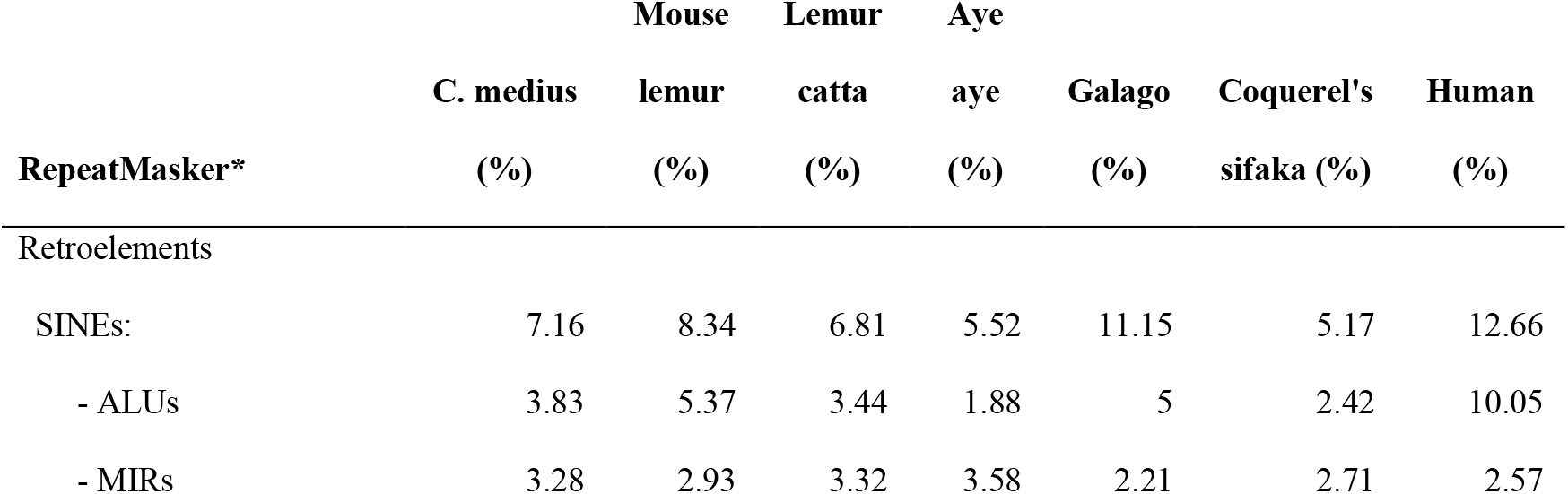
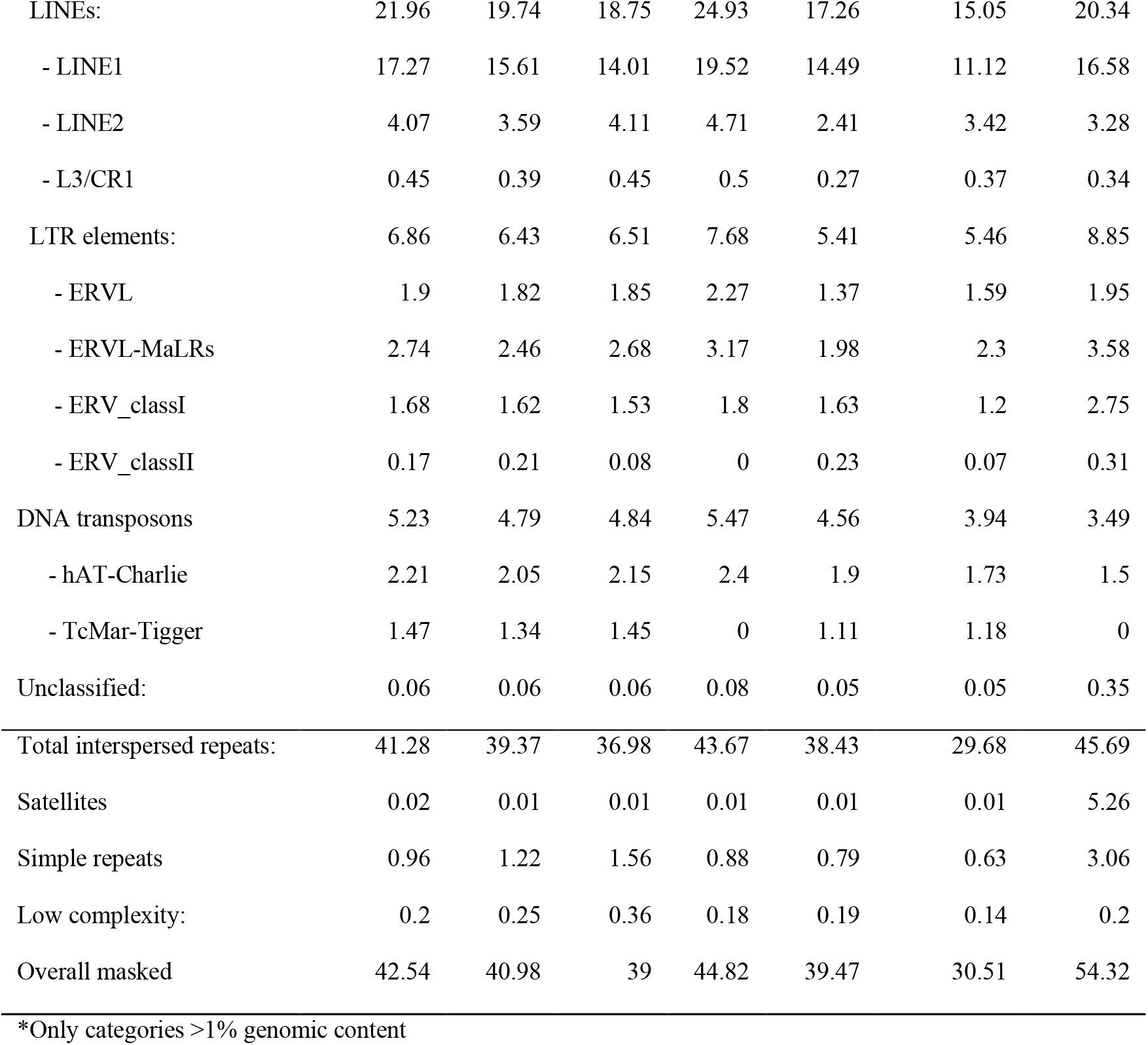
Complete repetitive DNA classification results for the *C. medius* genome assembly, FatTail1, generated with RepeatMasker, as compared to reference genomes of the other listed species generated with RepeatMasker.

### Heterozygosity and Demography

Tanager’s genome-wide heterozygosity index was 0.082%, calculated from dividing the total number of autosomal variants by the length of the genome (Table 5). There were a total of 1,892,877 Single Nucleotide Variants (SNVs) and 259,016 indels found by the combination of Clair3 and CuteSV. Only 41 variants were greater than 50 bp in length, the threshold of a structural variant. Ostrich’s heterozygosity index was 0.075% with 1,560,529 SNVs and 198,151 indel found. Both Tanager and Ostrich exhibited relatively low overall heterozygosity relative to previously published wild-caught *C. medius* individuals (0.1%)[33], consistent with historical bottlenecks, reduced long-term effective population size, and limited genetic diversity.

**Table 5:** Overall heterozygosity and ROH.

| Overall Heterozygosity | Tanager | Ostrich |
| --- | --- | --- |
|  | 0.082% | 0.075% |
| Runs of Homozygosity* |  |  |
| Total ROHs: | 15 | 2 |
| Total ROH length (Mb): | 163.71 | 6.00 |
| Mean ROH length (Mb): | 10.91 | 3.00 |
| Median ROH length (Mb): | 5.25 | 3.00 |
| Min ROH length (Mb): | 1.06 | 1.54 |
| Max ROH length (Mb): | 45.58 | 4.464 |
| F<sub>ROH</sub> (%): | 7.08% | 0.26% |

\*ROH stats (Filtered $\geq 1$ Mb)

Despite their similar overall heterozygosity, Tanager’s total length of runs of homozygosity (ROH) was dramatically larger than Ostrich. Tanager’s frequency of runs of homozygosity (FROH), the proportion of the genome that is homozygous above a 1 Mb length, was over 7% as compared to just 0.26% in Ostrich, suggesting strong recent inbreeding in Tanager specifically. Upon investigation, Tanager’s pedigree revealed a strong likelihood that his paternal grandfather was also his maternal uncle in addition to increased consanguinity in prior generations (Supplementary File 3). Ostrich is Tanager’s cousin twice removed, and the intervening 2 generations with outbred mating substantially reduced both the ROH and FROH to near levels seen in unrelated individuals while Tanager’s FROH is indicative of more recent inbreeding events that have not yet been fragmented by recombination. The lower heterozygosity observed in our captive individuals may partially reflect founder effects and recent breeding history within managed populations.

Effective population size was estimated for *C. medius* by the pairwise sequentially Markovian coalescent (PSMC) method (Figure 4). Population size history inferences on genomic runs of homozygosity were conducted with PSMC (Supplementary File 4). Despite differences in heterozygosity between the two individuals, the inferred demographic trajectories were broadly consistent across both genomes and supported a long-term decline in effective population size. The results suggested an effective population size (N_e_) near 5,000, with two major drops starting at 100,000 and 1 million years ago. Our estimates of N_e_ and historical demographics line up with the previous estimate of 98,005 reported by Williams et al. (2020), potentially reflecting differences between captive and wild demographic histories[33]. Our captive genomes preserved signatures of both historical demographic decline and recent managed breeding.

**Figure 4:**
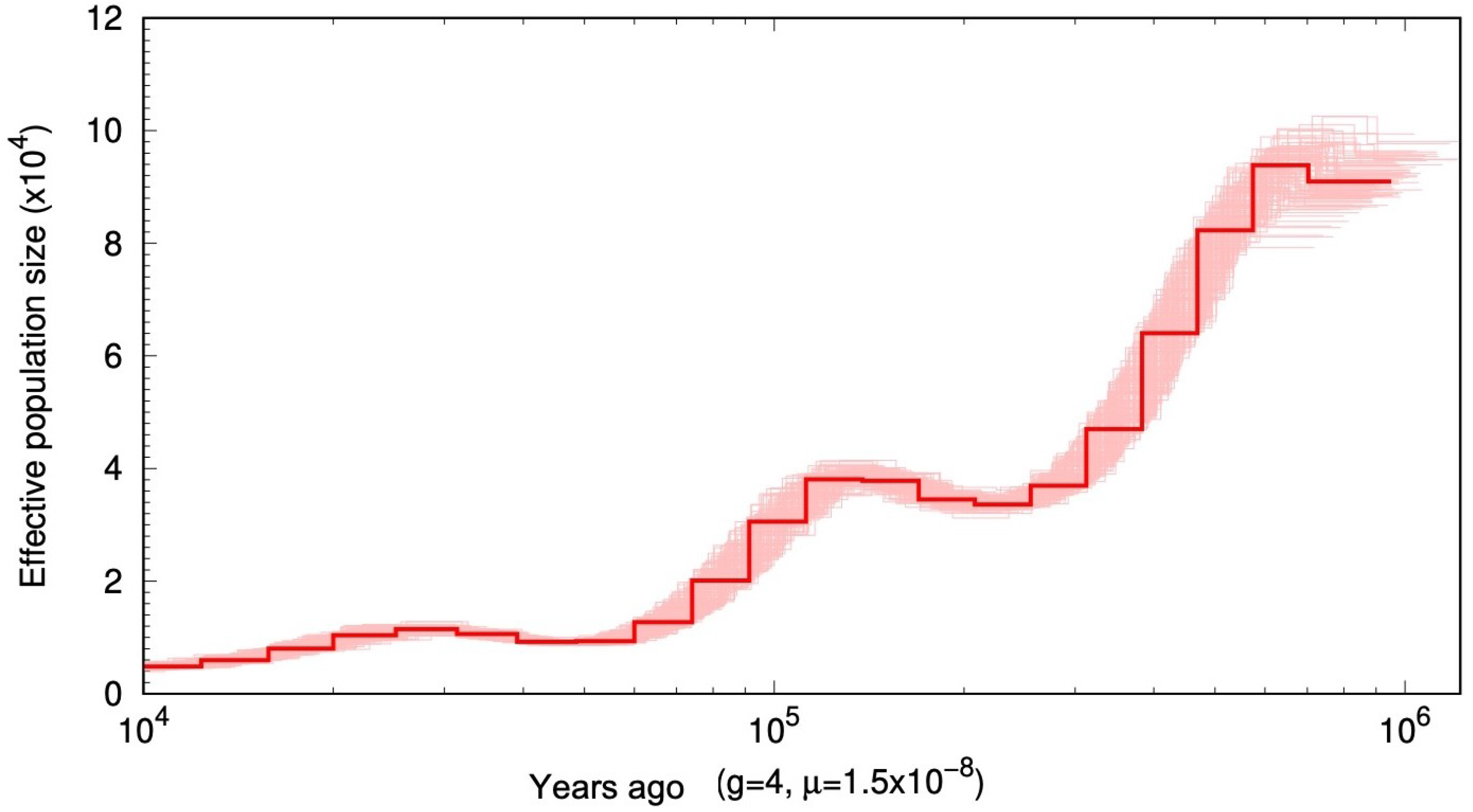
Effective population size generated with PSMC for the last 1 million years calculated with Tanager’s pseudohaplotypes. The red line represents the inferred demographic history, while lighter red lines indicate 100 bootstrap replicates. Generation time was estimated to be 4 years of age. Alt text: Pairwise Sequentially Markovian Coalescent (PSMC) plot showing inferred historical effective population size of *Cheirogaleus medius* over approximately one million years. The x-axis represents time before present on a logarithmic scale, and the y-axis shows effective population size. The red line represents the inferred demographic history, while lighter red lines indicate 100 bootstrap replicates. Effective population size increased over time with two major historical population expansions separated by periods of reduced population size.

Historical fluctuations in effective population size may be due to climatic changes during the Pleistocene, when glacial expansion caused the aridification of lowland areas[43]. The contractions of lowland forest habitat may have reduced population connectivity and reduced population size. Conversely, during warmer interglacial periods, suitable habitat increases would have coincided with increases in effective population size. Direct comparisons between our samples and those of Williams et al. (2020) should be interpreted cautiously due to differences in population origin, demographic history, and potential methodological variation. Williams et al. (2020) sampled an individual from an isolated population in sub-humid rainforest Tsihomanaomby in the north-east of Madagascar, outside of the typical habitat of *C. medius* of dry forests of western Madagascar. However, both studies see similar population expansions and contractions, concluding that *C. medius* has not recovered from the most recent population decline.

### Annotation

Homology-based gene annotation was performed with the NCBI tool Eukaryotic Genome Annotation Pipeline for external use (EGAPx) and uploaded to NCBI. Overall, 26,401 mRNAs were annotated, along with 23,925 genes, 686 non-coding RNAs, and 3991 pseudo-transcripts. Annotations were rated with a 99.17% completeness score via BUSCO in protein mode with the highest total and single copy score among high quality lemur genomes (Table 6). Despite the close relationship of lemurs to humans, annotation in this group has been poor. Even as recently as 2024, no annotation had existed for the aye-aye (*D. madagascariensis*) and the current annotation is still missing nearly 40% of genes[41]. Other lemur genomes, while better, are still lacking in annotation quality as compared to FatTail1, illustrating the improvement of both the genome and the EGAPx pipeline. However, an annotated chromosomal-level genome has now been published for the red-fronted brown lemur (*Eulemur rufifrons*) using the same EGAPx pipeline which allows direct end-user uploads to NCBI along with the genome[44]. Summary information describing the annotations is in Supplementary File 1.

**Table 6.**
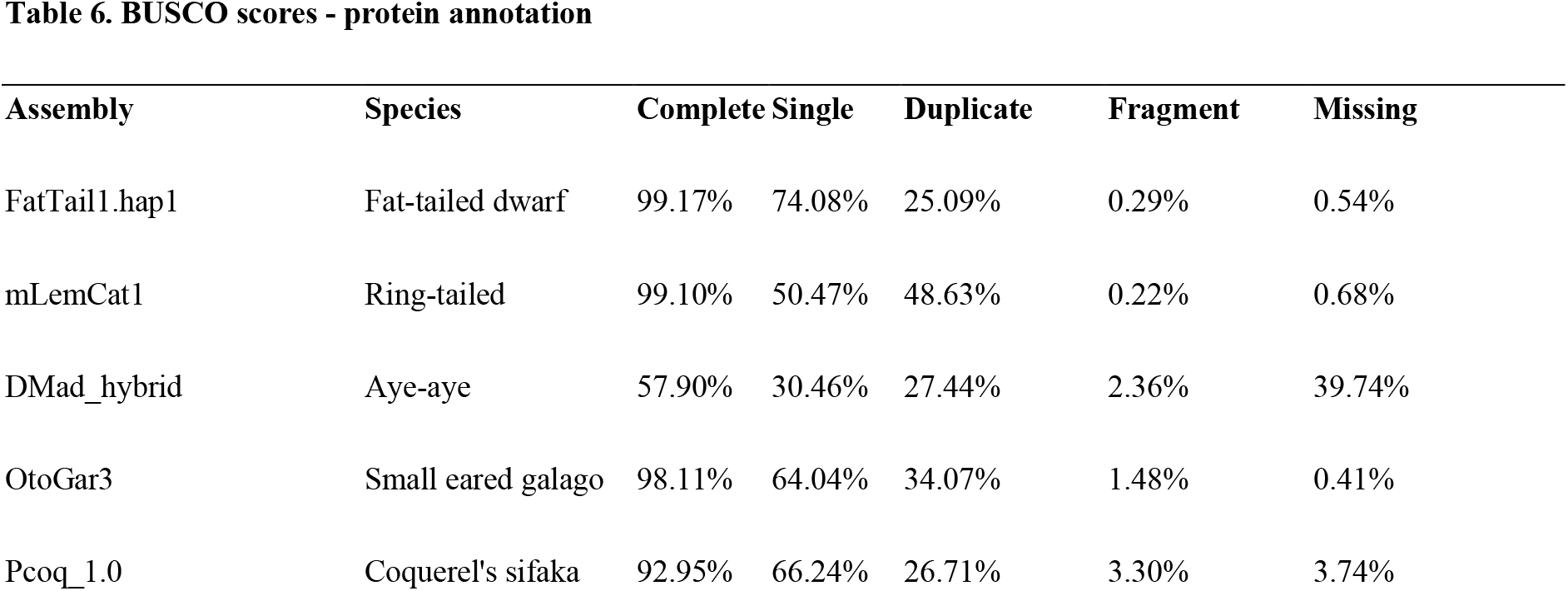
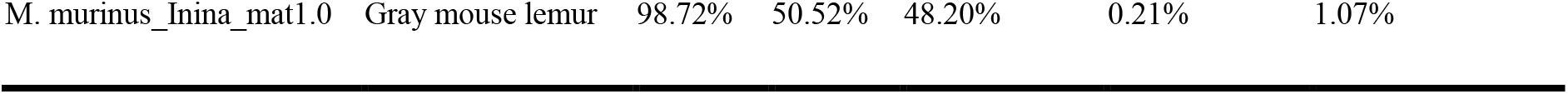
BUSCO scores - protein annotation.

### DNA Methylation

Nanopore instruments can detect native DNA modifications via modified pore signals compared to the unmodified base. Here, 5-methylcytosine (5mC) base modifications were called at CpG sites with FatTail1.hap1 as an alignment reference, generating methylation data for 25,867,291 CpG dinucleotides. Similar to blood samples in other primates, global DNA methylation at CpG dinucleotides was high at 65.21%.

Allele-specific methylation, a mechanism that facilitates allele-specific gene expression in genomic imprinting, was assessed by calling differentially methylated regions (DMRs) between pseudohaplotypes with DSS, representing each parent within Tanager’s genome. Strict calling parameters (i.e., methylation delta at least 65%, 100 bp length, at least 15 CpGs) yielded 37 unique loci (Supplementary File 1). Visualization of haplotagged read alignments at each DMR facilitated the discovery of 2 false positive loci, where apparent read misalignment for one of the two haplotypes led to a called DMR via a lack of CpGs on one allele. In total, 35 were deemed true DMRs following manual inspection. The validated DMRs had a mean length of 815 bp and a mean of 118 CpGs. A representative region is visualized in Figure 5 and is homologous to the known human imprinted region at PEG3/MEST.

**Figure 5:**
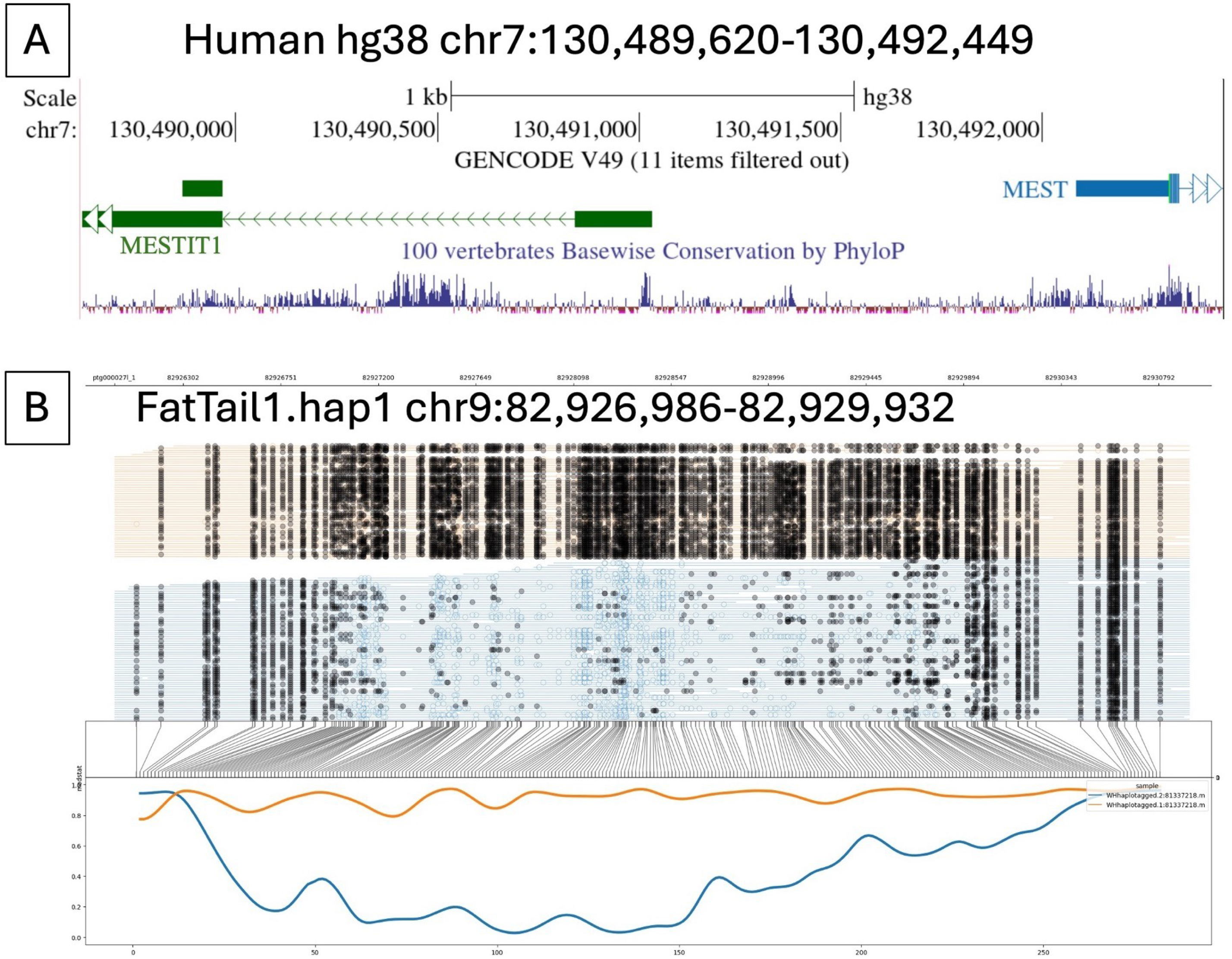
Allele-specific DNA methylation at a representative differentially methylated locus (DMR) called between *C. medius* assembly pseudohaplotypes. (A) Homologous region shown in human genome (hg38) maps this DMR to the known human imprinted gene, PEG1/MEST. (B) Reads mapping to each parental haplotype are shown in orange and blue respectively. Vertical lines indicate CpG positions within this 2 kb region. Open circles indicate unmethylated CpGs and black filled circles indicate methylated CpGs. In this region, nearly all methylated reads are associated with the haplotype 1 and unmethylated reads are associated with haplotype 2, despite there being no difference in underlying sequence. The flanking CpG positions show no parental difference. Alt text: Example of allele-specific DNA methylation at the MEST locus. Panel A shows the homologous human genomic region with gene annotations and evolutionary conservation. Panel B displays nanopore sequencing reads aligned to the corresponding region in the FatTail1 haplotype assembly, illustrating phased methylation patterns across individual reads. The lower plot shows methylation levels for the two parental haplotypes, demonstrating differential methylation consistent with allele-specific epigenetic regulation.

This method identifies predominantly imprinted gene loci in other species, though the strict parameters mean that many DMRs will be missed[45]. The resultant whole genome methylation data provides a baseline for studies involving environmentally sensitive changes to the epigenome including by aging, nutrition, stress, and other exposures[46].

### Mitochondrial Genome

The mitochondrial genome assembly built using MitoHiFi contained one circular contig 16,613 bp in length with 3000x read coverage. NCBI Blast revealed a 99.01% nucleotide identity match to a previously deposited *C. medius* mitogenome along with matches to additional *C. medius* mitogenomes down to 92.38%. The closest matches outside the species were to *C. major* and *C. crossleyi*, both at 88%.

Using our mitogenome and 9 others found on NCBI, a new phylogeny was built using *M. murinus* as the outgroup (Figure 6). The resulting phylogeny places our lemurs within the clade of other verified fat-tailed dwarf lemur mitogenomes. The mitogenomes of both Tanager and Ostrich cluster together and share 99.61% identity indicating very close kinship. The mitogenome was deposited as part of the scaffolded primary assembly at BioProject (Tanager PRJNA1484608 PRJNA1484609; Ostrich PRJNA1484630 PRJNA1484631).

**Figure 6:**
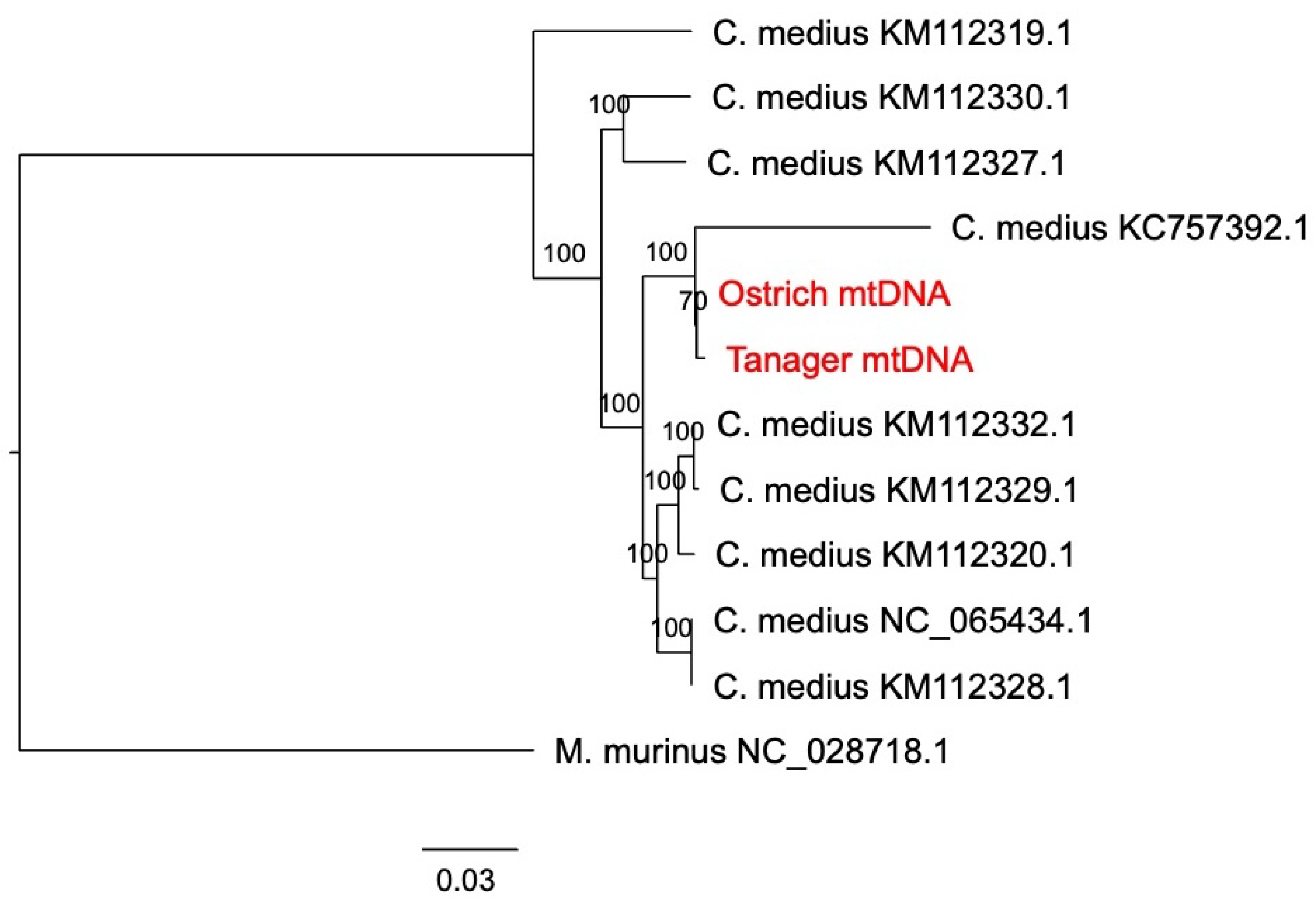
Phylogenetic relationship of Tanager and Ostrich’s mitogenomes derived through bootstrapped maximum likelihood comparison of the assembled mitogenome to 9 other complete *C. medius* mitogenomes available on NCBI. The tree is rooted on mouse lemur as the outgroup. Alt text: Phylogenetic tree depicting mitogenomes of 12 *C. medius*, and mouse lemur as the outgroup.

Mitochondrial divergence of ∼8% across publicly available *C. medius* mitogenomes substantially exceeds typical intraspecific values for mammals (generally <1.5% for cyt *b*)[47] and is consistent with the deep phylogeographic structure documented in this species complex[48,49]. This likely reflects the inclusion of geographically and evolutionarily distinct lineages, some of which have since been elevated to candidate or full species status, rather than variation within a single panmictic population. However, caution is warranted in using mitochondrial divergence alone as a species criterion, as mtDNA can overestimate divergence relative to the nuclear background, particularly in small mammals with variable substitution rates[50]. Given that there are reportedly large, heritable differences in body size, coloration, and other phenotypes in *C. medius* seen across Madagascar, species determination would benefit from a more comprehensive survey of *C. medius* across the island.

## DISCUSSION

We generated a chromosome-complete diploid genome assembly, mitogenome, and allele-specific methylation dataset for the fat-tailed dwarf lemur (*Cheirogaleus medius*) using nanopore sequencing. As the closest living relative to humans capable of prolonged hibernation, *C. medius* provides a unique comparative model for primate physiology, adaptation, and conservation genomics[1,33]. Prior strepsirrhine reference genomes, including those of the ring-tailed lemur (*L. catta*)[39], gray mouse lemur (*M. murinus*)[40], and aye-aye (*D. madagascariensis*)[41], substantially advanced comparative primate genomics. The markedly improved contiguity of our assembly enables higher-resolution analyses of structural variation, methylation, repetitive regions, and haplotype-specific genomic architecture. This assembly represents the most contiguous strepsirrhine genome produced to date prior to scaffolding and among the most contiguous nanopore-only strepsirrhine genome assemblies currently available, with our final genome having 38 contigs with an N50 of 103Mb, L50 of 10, and BUSCO of over 99%.

## POTENTIAL IMPLICATIONS

Our telomere-to-telomere reference genome of the hibernating fat-tailed dwarf lemur is a high-quality resource with broad use across multiple research fields. Using Oxford Nanopore long-read sequencing, we have substantially improved the foundation available for investigating the physiology and evolution of the fat-tailed dwarf lemur. As the only known obligately hibernating primate and closest living relative to humans capable of prolonged hibernation[1,2], *C. medius* provides a unique comparative model for understanding the molecular mechanisms that permit profound seasonal changed is metabolism and physiology. The high contiguity of FatTail1 will improve the accuracy of future transcriptomic[51,52], epigenomic[53], and comparative genomic analyses[54] and facilitate investigation of genomic regions that were difficult to resolve using previous, more fragmented assemblies[55,56].

Our combination of a highly contiguous genome assembly and DNA methylation data further demonstrates the utility of Oxford Nanopore sequencing for both characterizing the genome sequence and the epigenetic variation in a non-model primate. The haplotype-resolved methylation data allow for the investigation of allele-specific regulation and may facilitate future research on the regulatory mechanisms of tropical, primate hibernation[12]. Long-read sequencing further improves comprehensive analyses of transposable elements[57], structural variation[56,58] and other genomic features that are hindered by earlier, fragmented reference assemblies[59].

In addition to hibernation physiology, FatTail1 further facilitates comparative studies of primate genome evolution. Strepsirrhine primates are an early-diverging lineage within primate radiation[60], but are relatively underrepresented among high-quality reference genomes. The improved continuity of our assembly will allow comparisons across strepsirrhines and other primates and may help resolve lineage-specific patterns of genome structure, repetitive element evolution, and genetic variation.

Finally, the genome of *C.medius* has applications in conservation research. Lemurs are among the most threatened groups of mammals, threatened by anthropogenic deforestation and hunting[29]. High-quality reference genomes improve the characterization of genetic diversity, runs of homozygosity, demographic history, and potentially deleterious variation[32,61,62]. Although the individuals analyzed here originate from a managed captive population and therefore do not directly represent genetic diversity across wild populations, FatTail1 provides a substantially improved reference against which future population genomic data can be analyzed. Together, these resources establish a foundation for studies spanning primate hibernation, genome evolution, epigenetic regulation, and conservation genomics.

## DATA AVAILABILITY

The complete primary and secondary haplotype assemblies for both *C. medius* Tanager and Ostrich were deposited as BioProject PRJNA1484608 PRJNA1484609 (Tanager) and PRJNA1484630 PRJNA1484631 (Ostrich) in fasta format.

## ACKNOWLEDGEMENTS

We thank Tanager and Ostrich for providing the blood samples used in this study. This is Duke Lemur Center publication # XXXX. All procedures involving animals were approved by the Institutional Animal Care and Use Committee of Duke University (Protocol number: A208-23-10) and comply with all local laws.

## CONTRIBUTIONS TO AUTHORSHIP

AB and CF conceived and designed the study, provided funding, analyzed the data, and wrote the paper. RT performed the PSMC analysis. EE provided tissue samples and funding. CB performed pedigree analysis. AM and PL interpreted phylogenetic data and provided background and funding. All authors provided edits and approved the manuscript. Portions of the manuscript were edited with the assistance of OpenAI to improve grammar, clarity, and readability. All scientific interpretations, analyses, results, and conclusions were developed, verified, and approved by the authors, who take full responsibility for the content of the manuscript.

